# Mass Cytometric and Transcriptomic Profiling of Epithelial-Mesenchymal Transitions in Human Mammary Cell Lines

**DOI:** 10.1101/2021.03.26.436976

**Authors:** Johanna Wagner, Markus Masek, Andrea Jacobs, Charlotte Soneson, Nicolas Damond, Natalie de Souza, Mark D. Robinson, Bernd Bodenmiller

## Abstract

Epithelial-mesenchymal transition (EMT) equips breast cancer cells for metastasis and treatment resistance. Inhibition and elimination of EMT-undergoing cells are therefore promising therapy approaches. However, detecting EMT-undergoing cells is challenging due to the intrinsic heterogeneity of cancer cells and the phenotypic diversity of EMT programs. Here, we profiled EMT transition phenotypes in four non-cancerous human mammary epithelial cell lines using a FACS surface marker screen, RNA sequencing, and mass cytometry. EMT was induced in the HMLE and MCF10A cell lines and in the HMLE-Twist-ER and HMLE-Snail-ER cell lines by chronic exposure to TGFβ1 or 4-hydroxytamoxifen, respectively. We observed a spectrum of EMT transition phenotypes in each cell line and the spectrum varied across the time course. Our data provide multiparametric insights at single-cell level into the phenotypic diversity of EMT at different time points and in four human cellular models. These insights are valuable to better understand the complexity of EMT, to compare EMT transitions between the cellular models used herein, and for the design of EMT time course experiments.

|  |  |
| --- | --- |
| Measurement(s) | Human mammary epithelial cell lines • epithelial-mesenchymal transition • single-cell analysis |
| Technology Type(s) | Mass cytometry • Flow cytometry • RNA sequencing • Cell culture |
| Factor Type(s) | Epithelial-mesenchymal transition induced by TGFβ1 and 4-hydroxytamoxifen |
| Sample Characteristic - Organism | Homo sapiens |

Mendeley Data: DOI: 10.17632/pt3gmyk5r2.1

ArrayExpress Data: Accession number E-MTAB-9365

## Background & Summary

The epithelial-mesenchymal transition (EMT) equips epithelial cells with migratory, survival, and plasticity properties upon loss of epithelial hallmark characteristics. Together with its reverse process, the mesenchymal-epithelial transition, EMT contributes to cancer metastasis, provides resistance to cell death and chemotherapy, confers stemness properties to cancer cells, and interferes with immunotherapy^1–3^. EMT inhibition and elimination of EMT-undergoing cells are therefore investigated as approaches for cancer therapy^4^. However, detecting cancer cells undergoing EMT is challenging due to the intrinsic heterogeneity of cancer cells and the phenotypic diversity of EMT programs^4^.

A hallmark characteristic of epithelial cells is adhesion to neighboring cells and to the basement membrane^1^. To prevent anchorage-independent growth, epithelial cells normally undergo anoikis upon neighbor or matrix detachment^5^. During EMT, normal adhesion complexes, e.g. involving E-Cadherin, epithelial cell adhesion molecule (EpCAM), and laminin receptor integrin α6β1 (CD49f/CD29), are dissolved and resistance to anoikis is established^6,7^. Concomitant cytoskeletal rearrangements break down the epithelial apico-basal orientation and induce a motile front-back polarity, which often includes a replacement of cytokeratins with Vimentin^8^. EMT can further confer stemness properties to epithelial cells^9,10^. Numerous signaling pathways can trigger EMT, including TGFβ1, Notch, Hedgehog, WNT, and hypoxia, and activate downstream transcriptional drivers such as Snail family zinc finger transcription factors (TF), Twist family BHLH TFs, zinc finger E-box binding homeobox TFs, and homeobox TF PRRX1^11^. Regulation of EMT occurs by integration of epigenetic, transcriptional, post-transcriptional, and protein stability controls^11,12^. Together, this shows that the phenotypes of EMT-undergoing cells are shaped by complex molecular circuitries.

EMT is increasingly viewed more as a phenotypic continuum with intermediate states and less as a shift between two discrete states, and the concepts of ‘partial EMT’ and ‘hybrid EMT’ phenotypes have been introduced^4,13^. A systems biology approach used gene expression profiles of four non-small cell lung cancer cell lines to detect three intermediate states termed ‘pre-EMT’, ‘metastable EMT’, and ‘epigenetically-fixed’^14^. Transcriptomics of cell lines and clinical samples of cancer was used to rank the resulting spectrum of EMT states, showing that only some were linked to poor survival^15^. However, identification of EMT-undergoing cells in metastatic cancer tissue is still often based on co-expression of a few epithelial and mesenchymal markers^16,17^. This can be misleading as several of the ‘mesenchymal’ markers, e.g. Vimentin, can also be expressed by non-malignant epithelial cells^18^. It remains an ongoing debate which markers and combination of markers are sufficient to distinguish EMT from other processes *in vitro* and *in vivo*^4,19^. In particular, there remains the need for a comprehensive analysis of EMT phenotypes at the protein level.

To address this need, we applied multiplex single-cell mass cytometry^20^ to four non-cancerous human mammary epithelial cell lines that serve as widely-used models of EMT. EMT was induced in the HMLE and MCF10A cell lines by chronic exposure to TGFβ1^16,21^ and in the HMLE-Twist-ER (HTER) and HMLE-Snail-ER (HSER) cell lines by treatment with 4-hydroxytamoxifen (4OHT)^9^. In the HTER and HSER cell lines, 4OHT treatment allows the induction of gene expression by murine Twist1 fused to a modified estrogen receptor (ER) or SNAIL1-ER fusion protein, respectively^9^. To design our mass cytometry antibody panel, we conducted a fluorescence-based surface protein screen in parallel with a transcriptome analysis at multiple time points of induced EMT. We observed alterations in the surface proteome of EMT-undergoing cells over time and detected distinct gene expression profiles of hybrid epithelial-mesenchymal states compared with epithelial and mesenchymal states. From these analyses, we extracted candidate markers for multiplex mass cytometry, which revealed complex phenotypic transitions in all four EMT models and little phenotypic overlap of EMT states between the cell lines. The data presented here can aid in characterizing the complexity and dynamics of EMT in these widely used *in vitro* models.

## Methods

### Material

A table listing the material used in this study can be found on Mendeley Data (Online-only Table 1)^25^.

### Cell lines

The MCF10A human mammary epithelial cell line was obtained from the American Type Culture Collection (ATCC) and cultured in DMEM F12 Ham medium (Sigma Aldrich) supplemented with 10 μg/ml human insulin (Sigma Aldrich), 20 ng/ml epidermal growth factor (EGF, Peprotech), 500 ng/ml hydrocortisone (Sigma Aldrich), 5% horse serum (Gibco), 100 ng/ml cholera toxin (Sigma Aldrich), and PenStrep (Gibco). The HMLE, HMLE-Twist-ER (HTER), and HMLE-Snail-ER (HSER) cell lines were a gift from the laboratory of Prof. Robert Weinberg at the Whitehead Institute for Biomedical Research and the Massachusetts Institute of Technology and were cultured in a 1:1 mixture of DMEM F12 Ham medium (Sigma Aldrich) supplemented with 10 μg/ml human insulin (Sigma Aldrich), 10 ng/ml EGF (Peprotech), 500 ng/ml hydrocortisone (Sigma Aldrich), and PenStrep (Gibco) with the mammary epithelial growth medium (MEGM^TM^) BulletKit^TM^ (Lonza). For the HTER and HSER cell lines, the growth medium was supplemented with 1 μg/ml Blasticidin S (InvivoGen).

### EMT time courses and cell harvesting

EMT was induced in the MCF10A cell line by chronic stimulation with 5 ng/ml TGFβ1 (Cell Signaling Technology) for eight days^22^. For this, 0.8 million cells were seeded per 10 cm cell culture dish (Nunc) and incubated at 37 °C and 5% CO_2_ according to ATCC recommendations. TGFβ1 treatment and vehicle treatment using Dulbecco’s phosphate buffer saline (PBS, Sigma Aldrich) started 24 hours after seeding and was applied daily together with a growth medium exchange.

EMT was induced in the HMLE cell line by chronic stimulation with 4 ng/ml TGFβ1 (Cell Signaling Technology) for 14 days^9^. For this, 0.5 million cells were seeded per 10 cm cell culture dish (Nunc) and incubated at 37 °C and 5% CO_2_. TGFβ1 treatment and vehicle treatment using PBS (Sigma Aldrich) started 24 hours after seeding and was applied daily. The growth medium was exchanged every other day.

EMT was induced in the HTER and HSER cell lines by chronic stimulation with 4 ng/ml 4-hydroxytamoxifen (4OHT; Sigma Aldrich) for 14 days^9^. For this, 0.5 million cells were seeded per 10 cm cell culture dish (Nunc) and incubated at 37 °C and 5% CO_2_. 4OHT treatment and vehicle treatment using methanol (Thommen Furler) started 24 hours after seeding and was applied daily. The growth medium was exchanged every other day.

To avoid over-confluence and senescence during the time course of HMLEs, HTERs, and HSERs, the cells were split and re-seeded on day four and eight. For this, the cells were washed once with pre-warmed PBS, incubated for 5 min at 37 °C with 4 ml pre-warmed TrypLE 1X Express (Gibco), quenched with pre-warmed growth medium, pelleted at 350 x g for 5 min at room temperature, resuspended in pre-warmed growth medium, and re-seeded using 0.5 million cells per 10 cm cell culture dish.

For harvesting, the cells were washed once with pre-warmed PBS, incubated for 5 min at 37 °C with pre-warmed TrypLE 1X Express (Gibco), fixed for 10 min at room temperature with 1.6% paraformaldehyde (PFA, Electron Microscopy Sciences), scraped off the dish using a cell scraper (Sarstedt AG), and quenched using 4 °C growth medium. The cells were pelleted at 600 x g for 4 min at 4 °C, resuspended in 4 °C PBS at a concentration of about 0.5 million cells per ml and frozen at −80 °C. For mass cytometry analysis, 5-Iodo-2′-deoxyuridine (IdU) at 10 μM was added to the medium 20 min before cell harvesting^23^.

### Mass-tag cellular barcoding

To minimize inter-sample staining variation, we applied mass-tag barcoding to fixed cells^24^. A barcoding scheme composed of unique combinations of four out of nine barcoding metals was used for this study; metals included palladium (^105^Pd, ^106^Pd, ^108^Pd, ^110^Pd, Fluidigm) conjugated to bromoacetamidobenzyl-EDTA (Dojindo) as well as indium (^113^In and ^115^In, Fluidigm), yttrium, rhodium, and bismuth (^89^Y, ^103^Rh, ^209^Bi, Sigma Aldrich) conjugated to maleimido-mono-amide-DOTA (Macrocyclics). The concentrations were adjusted to 20 nM (^209^Bi), 100 nM (^105-110^Pd, ^115^In, ^89^Y), 200 nM (^113^In), or 2 μM (^103^Rh). Cells were randomly distributed across a 96-well plate and about 0.3 million cells per well were barcoded using a transient partial permeabilization protocol. Cells were washed once with 0.03% saponin in PBS (Sigma Aldrich) prior to incubation in 200 μl barcoding reagent for 30 min at room temperature. Cells were then washed four times with cell staining medium (CSM, PBS with 0.3% saponin, 0.5% bovine serum albumin (BSA, Sigma Aldrich) supplemented with 2 mM EDTA (Stemcell Technologies) and pooled for antibody staining.

### Fluorescence cellular barcoding and FACS surface protein screen

To apply the FACS surface protein screen to multiple samples simultaneously, we performed fluorescence barcoding of fixed cells. For this, 18 million cells were washed once with CSM prior to incubation in 3 ml barcoding reagent for 20 min at 4 °C in the dark. As barcoding reagents Alexa Fluor-700-NHS-Ester (AF700, Molecular Probes) and Pacific Orange-NHS-Ester (PO, Molecular Probes) dissolved in dimethyl sulfoxide (DMSO) at 200 μg/ml were used. Single stains or a combination of AF700 and PO were performed in CSM at a final concentration of 0.1 μg/ml or 1 μg/ml and 0.4 μg/ml or 2 μg/ml, respectively. Cells were washed twice with CSM before pooling and staining with E-Cadherin-AF647 (clone 67A4, Biolegend) and EpCAM-FITC (clone 9C4, Biolegend) or CD44-FITC (clone IM7, Biolegend) for 20 min at 4 °C in the dark. Cells were washed once with CSM and filtered through a 40 μm cell strainer. About 0.3 million cells in 37.5 μl CSM were loaded in each well of a 96-well plate of the Human Cell Surface Marker Screening (phycoerythrin [PE]) Kit (Biolegend). Each well contained 12.5 μl of diluted PE-conjugated antibody in CSM. The cells were incubated for 30 min at 4 °C in the dark, according to manufacturer’s instructions. The cells were then washed twice with CSM, fixed with 1.6% PFA in PBS for 10 min at room temperature in the dark and washed twice with CSM again, prior to FACS analysis using the LSRFortessa Cell Analyzer (BD Biosciences).

### FACS sorting and RNA sequencing

For live cell FACS sorting, cells were washed once with pre-warmed PBS, incubated for 5 min at 37 °C with 4 ml pre-warmed TrypLE 1X Express (Gibco), pipetted off the cell culture dish, and collected in 4 °C PBS. Cells were pelleted at 350 x g for 5 min at 4 °C, re-suspended in 4 °C PBS with 1% BSA, and stained with E-Cadherin-AF647 (clone 67A4, 5 μg/ 100 μl, Biolegend) and CD44-PE (clone IM7, 1.25 μg/ 100 μl, Biolegend) for 20 min at 4 °C in the dark. Cells were washed once using PBS with 1% BSA and kept on ice until FACS sorting using the FACSAria III (BD Biosciences). For RNA isolation, cells were pelleted at 350 x g for 5 min at 4 °C and lysed in 350 μl RLT buffer of the RNeasy Mini Kit (Qiagen). RNA was isolated according to the manufacturer’s instructions. Briefly, RNA was collected on the RNeasy spin column, washed with 70% ethanol (Merck), and DNA was removed by incubation with DNAse I (Qiagen). RNA was collected in 30-50 μl diethylpyrocarbonate (DEPC, Sigma Aldrich)-containing water and stored at −80 °C. DEPC water was prepared by dissolving 1 ml DEPC in 1 L ddH_2_O prior to autoclaving. The RNA quality was assessed using a NanoDrop (Thermo Scientific) and Bioanalyzer (Agilent). RNA sequencing was performed using the HiSeq 2500 System (Illumina) in SR 50 mode (50 base reads) after poly (A) enrichment and stranded library preparation.

### Antibodies and antibody labeling

All antibodies and corresponding clone, provider, and metal or fluorescence tag are listed in the Online-only Table 1 and Online-only Table 20 on Mendeley Data^25^. Target specificity of the antibodies was confirmed in our laboratory. Antibodies were obtained in carrier/ protein-free buffer or were purified using the Magne Protein A or G Beads (Promega) according to manufacturer’s instructions. Metal-labeled antibodies were prepared using the Maxpar X8 Multimetal Labeling Kit (Fluidigm) according to manufacturer’s instructions. After conjugation, the protein concentration was determined using a NanoDrop (Thermo Scientific), and the metal-labeled antibodies were diluted in Antibody stabilizer PBS (Candor Bioscience) to a concentration of 200 or 300 μg/ml for long-term storage at 4 °C. Optimal concentrations for antibodies were determined by titration, and antibodies were managed using the cloud-based platform AirLab as previously described^26^.

### Antibody staining and cell volume quantification

Antibody staining was performed on pooled samples after mass-tag cellular barcoding. The pooled samples were washed once with CSM. For staining with the EMT antibody panel (Online-only Table 20 on Mendeley Data^25^), cells were incubated for 45 min at 4 °C followed by three washes with CSM. For mass-based cell detection, cells were stained with 500 μM nucleic acid intercalator iridium (^191^Ir and ^193^Ir, Fluidigm) in PBS with 1.6% PFA (Electron Microscopy Sciences) for 1 h at room temperature or overnight at 4 °C. Cells were washed once with CSM and once with 0.03% saponin in PBS. For cell volume quantification, cells were stained with 12.5 μg/ml Bis(2,2′-bipyridine)-4′-methyl-4-carboxybipyridine-ruthenium-N-succidimyl ester-bis(hexafluorophos-phate) (^96^Ru, ^98-102^Ru, ^104^Ru, Sigma Aldrich) in 0.1 M sodium hydrogen carbonate (Sigma Aldrich) for 10 min at room temperature as previously described^23^. Cells were then washed twice with CSM, twice with 0.03% saponin in PBS, and twice with ddH_2_O. For mass cytometry acquisition, cells were diluted to 0.5 million cells/ml in ddH_2_O containing 10% EQ^TM^ Four Element Calibration Beads (Fluidigm) and filtered through a 40 μm filter cap FACS tube. Samples were placed on ice and introduced into the Helios upgraded CyTOF2 (Fluidigm) using the Super Sampler (Victorian Airship) introduction system; data were collected as .fcs files.

## Statistical Analysis

### Mass cytometry data preprocessing

Mass cytometry data were concatenated using the .fcs File Concatenation Tool (Cytobank, Inc.), normalized using the MATLAB version of the Normalizer tool^27^, and debarcoded using the CATALYST R/Bioconductor package^28^. The .fcs files were uploaded to the Cytobank server (Cytobank, Inc.) for manual gating on populations of interest. The resulting population was exported as .fcs files and loaded into R (R Development Core Team, 2015) for downstream analysis.

### FACS surface marker screen data processing

FACS data were compensated on the LSRFortessa Cell Analyzer (BD Biosciences) using single-stained samples. The .fcs files were uploaded to the Cytobank server (Cytobank, Inc.) for manual debarcoding and gating on populations of interest. The mean signal intensity per well and population of interest was exported as an excel sheet. The mean signal intensity of the ‘Blank’ wells of the screen and the signal intensity of the respective ‘Isotype control’ well were subtracted. From the resulting intensity values, log2-transformed fold changes were calculated.

### Dimensionality reduction analyses

For dimensionality reduction visualizations using the t-SNE and UMAP algorithms^29,30,48^, signal intensities (dual counts) per channel were arcsinh-transformed with a cofactor of 5 (counts_transf = asinh(x/5)). The R t-SNE package for Barnes-Hut implementation and the R UMAP implementation package *uwot* (https://github.com/jlmelville/uwot) were used. For marker expression level visualization on t-SNE plots, the expression was normalized between 0 and 1 to the 99^th^ percentile and the top percentile was set to 1.

### RNA sequencing data analysis

The RNA sequencing data was processed using an analysis setup derived from the ARMOR workflow^31^. Quality control of the raw FASTQ files was performed using FastQC v0.11.8 (Andrews S, Babraham Bioinformatics, https://www.bioinformatics.babraham.ac.uk/projects/fastqc/). Transcript abundances were estimated using Salmon v1.2.0^32^, using a transcriptome index based on Gencode release 34^33^, including the full genome as decoy sequences^34^ and setting the k-mer length to 23. For comparison, the reads were also aligned to the genome (GRCh38.p13) using STAR v2.7.3a^35^. Transcript abundances from Salmon were imported into R v4.0.2 and aggregated on the gene level using the tximeta Bioconductor package, v1.6.2^36^. The quasi-likelihood framework of edgeR, v3.30.0^37,38^ was used to perform differential gene expression analysis, accounting for differences in the average length of expressed transcripts between samples^39^. In each comparison, edgeR was used to test the null hypothesis that the true absolute log2-fold change between the compared groups was less than 1. edgeR was also used to perform exploratory analysis and generate a low-dimensional representation of the samples using multidimensional scaling (MDS). The analysis scripts were run via Snakemake^40^, and all the code is available on GitHub^41^.

## Data Records

A detailed list of all materials used in this study can be found as Online-only Table 1 on Mendeley Data^25^. RNA sequencing data have been deposited in the ArrayExpress database at EMBL-EBI with accession number E-MTAB-9365^42^. Tables showing the results of the differential gene expression analyses and a table reporting the RNA quality and RNA sequencing mapping metrics have been deposited as Online-only Tables 2-13 on Mendeley Data^25^. The code used for RNA sequencing data analysis can be found on GitHub^41^. FACS surface protein screen data as .fcs files and the corresponding data analyses referenced in the text as Online-only Tables 14-19 have been deposited on Mendeley Data^25^. Furthermore, the Biolegend data sheet corresponding to the FACS screen has been deposited^25^. Mass cytometry .fcs files of cells after debarcoding (‘DebarcodedCellsGate’) and of live cells (‘LiveCellsGate’) have been deposited on Mendeley Data^25^ together with a table containing .fcs file annotations (‘FCS_File_Information’) and a table corresponding to the antibody panel used (Online-only Table 20).

## Technical Validation

### Optimizing the time courses for *in vitro* induction of EMT

We induced EMT in four non-cancerous human mammary epithelial cell lines by chronic ectopic stimulation with TGFβ1 or 4OHT over several days (Figure 1a; Methods); all four systems are widely used models of EMT^9,16,21^. We initially carried out a basic characterization of these models and optimized each induction time course to yield the maximum percentage of cells with mesenchymal (M) phenotype, characterized by loss of E-Cadherin and concomitant gain of expression of Vimentin^4^. We excluded apoptotic cells from the analysis (Figure 1b).

**Figure 1.**
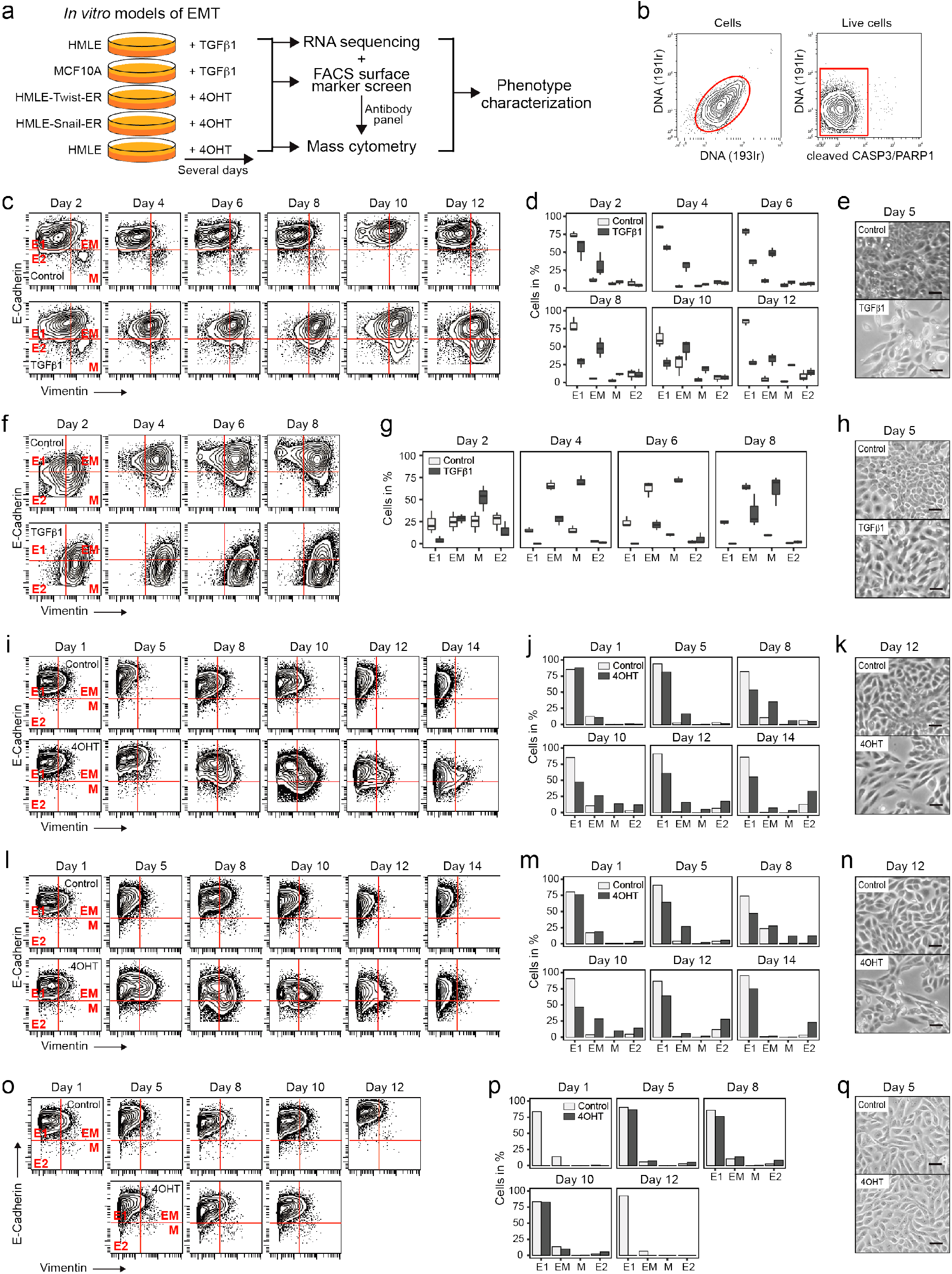
Induction of EMT in human mammary epithelial cell lines. (a) Experimental workflow. (b) Gating to select live cells. (c) E-Cadherin and Vimentin expression in HMLEs. Gating to select populations with E1-, E2-, EM-, or M-phenotype. (d) Percentages of HMLEs per gate and time point as in (c). (e) Phase contrast images of HMLEs. (f) E-Cadherin and Vimentin expression in MCF10As. (g) Percentage of MCF10As cells per gate and time point as in (f). (h) Phase contrast images of MCF10As. (i) E-Cadherin and Vimentin expression in HTERs. (j) Percentage of HTERs per gate and time point as in (i). (k) Phase contrast images of HTERs. (l) E-Cadherin and Vimentin expression in HSERs. (m) Percentage of HSERs per gate and time point as in (l). (n) Phase contrast images of HSERs. (o) E-Cadherin and Vimentin expression in HMLEs. (p) Percentage of HMLEs per gate and time point as in (o). (q) Phase contrast images of HMLEs. Scale bar = 10 μm. E1 = epithelial 1, E2 = epithelial 2, EM = hybrid epithelial-mesenchymal, M = mesenchymal.

On day 12 of chronic exposure to TGFβ1, the HMLE cell line yielded 25% of cells with an M-phenotype, 33% of cells with a hybrid epithelial-mesenchymal (EM) phenotype with increased Vimentin expression but no downregulation of E-Cadherin, 28% of an E-Cadherin^high^Vimentin^low^ phenotype (E1), and 14% of an E-Cadherin^low^Vimentin^low^ phenotype (E2) (Figures 1c and 1d). In comparison, on day twelve, 2% of control HMLEs exhibited an M-phenotype, 5% an EM-phenotype, 84% an E1-phenotype, and 9% an E2-phenotype (Figures 1c and 1d). Control HMLEs with EM- or E2-phenotype were most abundant during sparse growth conditions, such as after splitting (Figure 1d, Methods), indicating a regulation of E-Cadherin and Vimentin levels by growth density^16,43^. As previously reported, treatment with TGFβ1 induced spindle-like morphological changes^44^ and resulted in lower cell density compared with control^45^ (Figure 1e).

In the MCF10A cell line, induction of EMT by TGFβ1 treatment occurred in a different time frame. The percentage of cells with an M-phenotype increased from 54% on day two to 70% on day eight, the percentage of EM cells (28%) and E1 cells (2%) remained stable across the time course, and the percentage of E2 cells dropped from 10% to 2% (Figures 1f and 1g). In control, cells with M-phenotype were at 26% on day 2 and 10% on day 8, cells with EM phenotype more than doubled from 25% to 64%, the percentage of E1 cells stayed stable at 22%, and the E2 cells decreased from 29% to 1% over the time course (Figures 1f and 1g). As reported, TGFβ1-treated MCF10A cells acquired spindle-like morphologies while control cells retained their cobblestone shape (Figure 1h)^16^. Together, these data show that under sparse growth conditions on day 2, MCF10A cells exhibit mesenchymal-like phenotypes even without TGFβ1 treatment, reflecting the basal-like character of the cell line^16^. An increase in cell density over time is accompanied by upregulation of E-Cadherin and therefore loss of the M-phenotype in control, while stimulation with TGFβ1 inhibits an E-Cadherin upregulation and induces an upregulation of Vimentin. In TGFβ1-treated cells, a decrease in the percentage of cells with M-phenotype on day eight compared with day six, suggests that cell density may inhibit further EMT^46^.

In the HTER and HSER cell lines, EMT was induced by chronic treatment with 4OHT (Methods). We detected the highest percentage (14%) of 4OHT-treated HTER cells with M-phenotype on day ten, at which point 26% of cells exhibited an EM-phenotype (Figures 1i and 1j). The percentage of 4OHT-treated HSER cells with M-phenotype peaked at 12% on day eight and 28% of cells exhibited an EM-phenotype at this time point (Figures 1l and 1m). For both cell lines, treatment with 4OHT induced spindle-like morphologies and was accompanied by reduced cell density compared with control (Figure 1k and 1n), as previously reported^9^. We then assessed possible effects of the 4OHT treatment on HMLEs in the absence of the Twist1-ER or SNAIL1-ER fusion proteins. As expected, treatment with 4OHT did not induce EMT or morphological changes in HMLEs (Figures 1o and 1p). In treated and control, the percentage of cells with M-phenotype was below 1% and cells with EM-phenotype at 11% at all time points, indicating a basal-like character of the cell line^9^. The majority of treated and control HMLEs maintained an E1-phenotype throughout the time course (Figure 1o).

In conclusion, we could induce EMT in four *in vitro* human cell line models of this process. We observed phenotypic variability, including both full and partial EMT phenotypes, in response to 1-2 weeks of chronic stimulation with TGFβ1 or 4OHT. Each model followed a unique EMT timeline and showed varying extents of transition to the mesenchymal phenotype.

### Transcriptomic profiling of cells undergoing EMT

We next used RNA sequencing to identify markers that distinguish EMT-undergoing cells from control and markers that distinguish cells with EM-phenotype from cells with E- or M-phenotype. From the resulting markers, candidates were selected to inform a mass cytometry antibody panel design. For RNA sequencing, EMT-undergoing HTER cells on day eight and day twelve were sorted by fluorescence-activated cell sorting (FACS) into three populations: E-Cadherin^high^CD44^low^ (E1-phenotype), E-Cadherin^int^CD44^int^ (EM-phenotype), and E-Cadherin^low^CD44^high^ (M-phenotype) (Figure 2a, Methods). CD44 served as a surrogate M-phenotype marker for intracellular Vimentin to avoid cell permeabilization and RNA loss^9^. As control, day-matched untreated HTER cells with E1-phenotype were used (Figure 2a). As a second type of control to monitor possible effects of 4OHT independent of EMT, we included 4OHT-treated and untreated HMLE cells. We included two to four pairs of independent biological replicates per condition and collected high quality RNA for all samples (Online-only Table 2, Methods).

**Figure 2.**
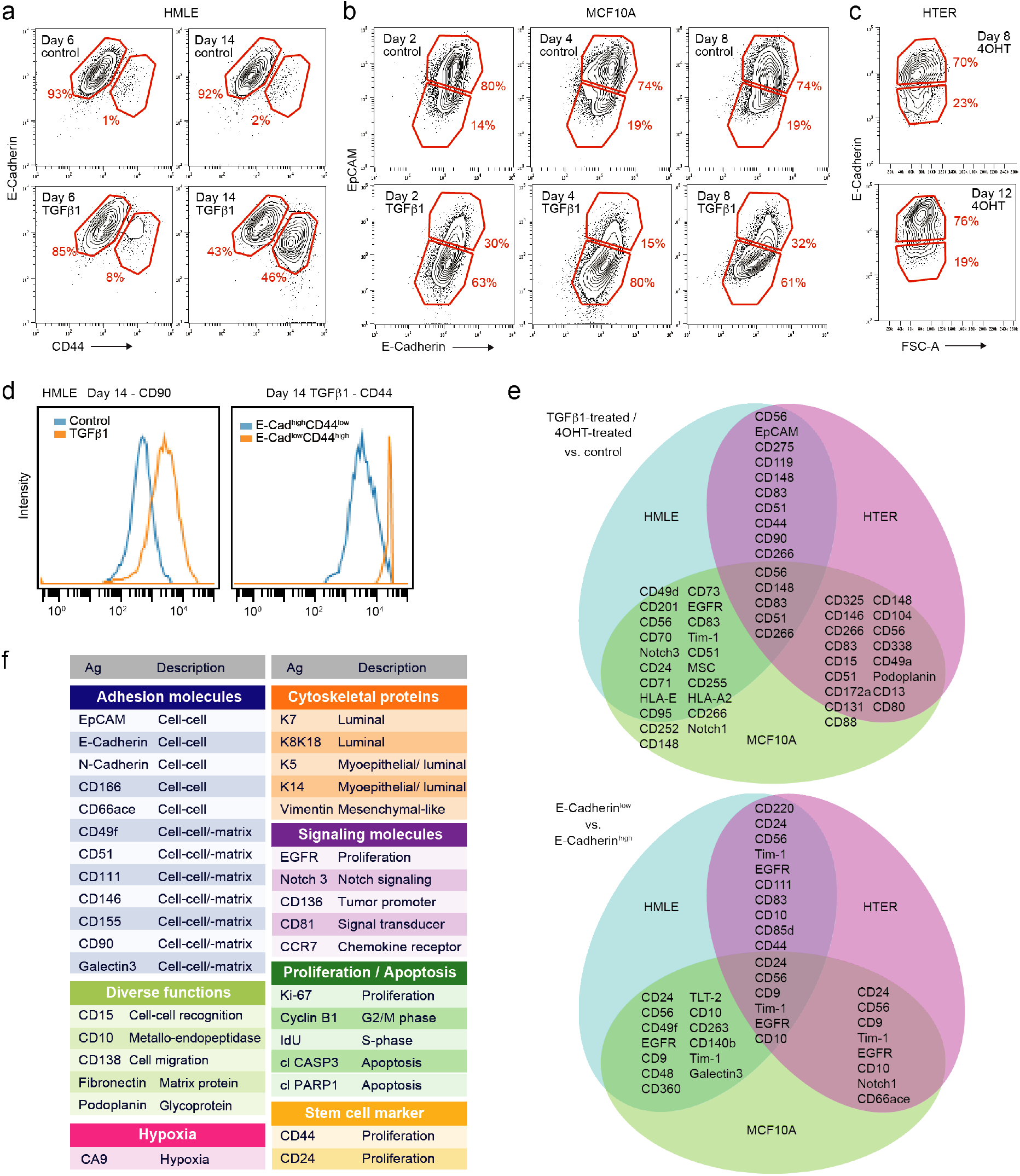
Transcriptomic profiling of EMT-undergoing mammary epithelial cells. (a) Gating to select populations of interest of HTERs for RNA sequencing. (b) Number of RNA sequencing reads assigned to genes per sample. (c) Average base quality (upper panel) and GC content (lower panel) for all samples. (d) Multidimensional scaling plot showing the first two dimensions. (e-g) Volcano plots showing the indicated differential gene expression analyses. Highlighted in red are genes with an adjusted p-value below 0.05. logFC = log2 fold change, E = epithelial, EM = hybrid epithelial-mesenchymal, M = mesenchymal.

RNA sequencing yielded above 20 million reads per sample assigned to genes, except one sample with 19 million reads (Figure 2b, Online-only Table 2). Mean Phred scores ranged between 35 and 36, indicating high base call accuracy, and GC content distribution across samples did not indicate any noticeable contamination (Figure 2c, Online-only Table 2). For all samples, more than 82% of the reads could be uniquely aligned to the human reference genome using STAR^35^. Mapping to the transcriptome index using Salmon^32^ showed that more than 86% of fragments were assigned to a transcript, with little variation across samples.

We next assessed the similarity of samples based on global gene expression levels using multidimensional scaling^37,38^ (Methods). This showed that the respective pairs of biological replicates were similar (Figure 2d). Control HTER cells were similar to day-matched 4OHT-treated and control HMLE cells, indicating few effects of 4OHT on transcription independent of EMT. This analysis further revealed that 4OHT-treated HTER cells with E-, EM-, and M-phenotype were all separate from their respective day-matched control (Figure 2d). Differential gene expression analysis showed that more genes were significantly differentially expressed between HTER cells with M-phenotype or EM-phenotype and control than between E-phenotype and control on day eight (Figure 2e, Online-only Tables 3–5). Among differentially expressed genes between M-phenotype and control, we found upregulation of canonical markers of EMT^1^, such as the transcription factors *ZEB1, ZEB2, FOXC2*, and *PRRX1*, as well as downregulation of typical epithelial markers such as *EPCAM* (Online-only Table 3). We then asked, which genes were significantly differentially expressed between HTER cells with EM-phenotype and cells with E- or M-phenotype on day eight and found three genes (*HHIP, FBN1, HHIP-AS1*) and one gene (*KIAA1755*), respectively (Figure 2f, Online-only Tables 6 and 7). When comparing HTER cells on day twelve, more genes were significantly differentially expressed between cells with M-phenotype and control than between E-phenotype and control (Figure 2g, Online-only Tables 8 and 9).

In conclusion, 4OHT-treated HTER cells with M-phenotype or EM-phenotype deviated transcriptionally more from control than cells with E-phenotype. Also, 4OHT-treated cells with E-phenotype are transcriptionally distinct from control cells with E-phenotype.

### Surface protein expression screen during EMT

We then carried out a FACS-based surface protein screen to identify further markers that distinguish EMT-undergoing cells from control and M-phenotype cells from E-phenotype cells, for design of the mass cytometry antibody panel. Treated and control samples of the HTER, HMLE, and MCF10A cell lines were fixed at multiple time points, fluorescently barcoded, and co-stained with a combination of surface epithelial markers, E-Cadherin and/or EpCAM, and a surface mesenchymal marker, CD44, to detect M-and E-phenotypes. The resulting FACS data were compensated, debarcoded and gated for cell populations of interest (Figures 3a-c, Methods). We detected expected surface protein abundance differences between cell populations, confirming the quality of the screening results (Figure 3d). We identified multiple surface proteins that were more than two-fold differentially expressed between treated (TGFβ1-treated or 4OHT-treated) and control samples (Tables 1–3, Online-only Tables 14-16), several of which (e.g., CD90, CD146, CD166, CD51, and Podoplanin) were regulated in more than one cell line (Figure 3e, upper panel). Similarly, we identified multiple surface proteins that were differentially expressed between cells with M-phenotype and cells with E-phenotype (Tables 4–6, Online-only Tables 17-19), several of which were again shared between cell lines, such as CD24, CD56, CD9, TIM-1, EGFR, and CD10 (Figure 3e, lower panel).

**Table 1:**
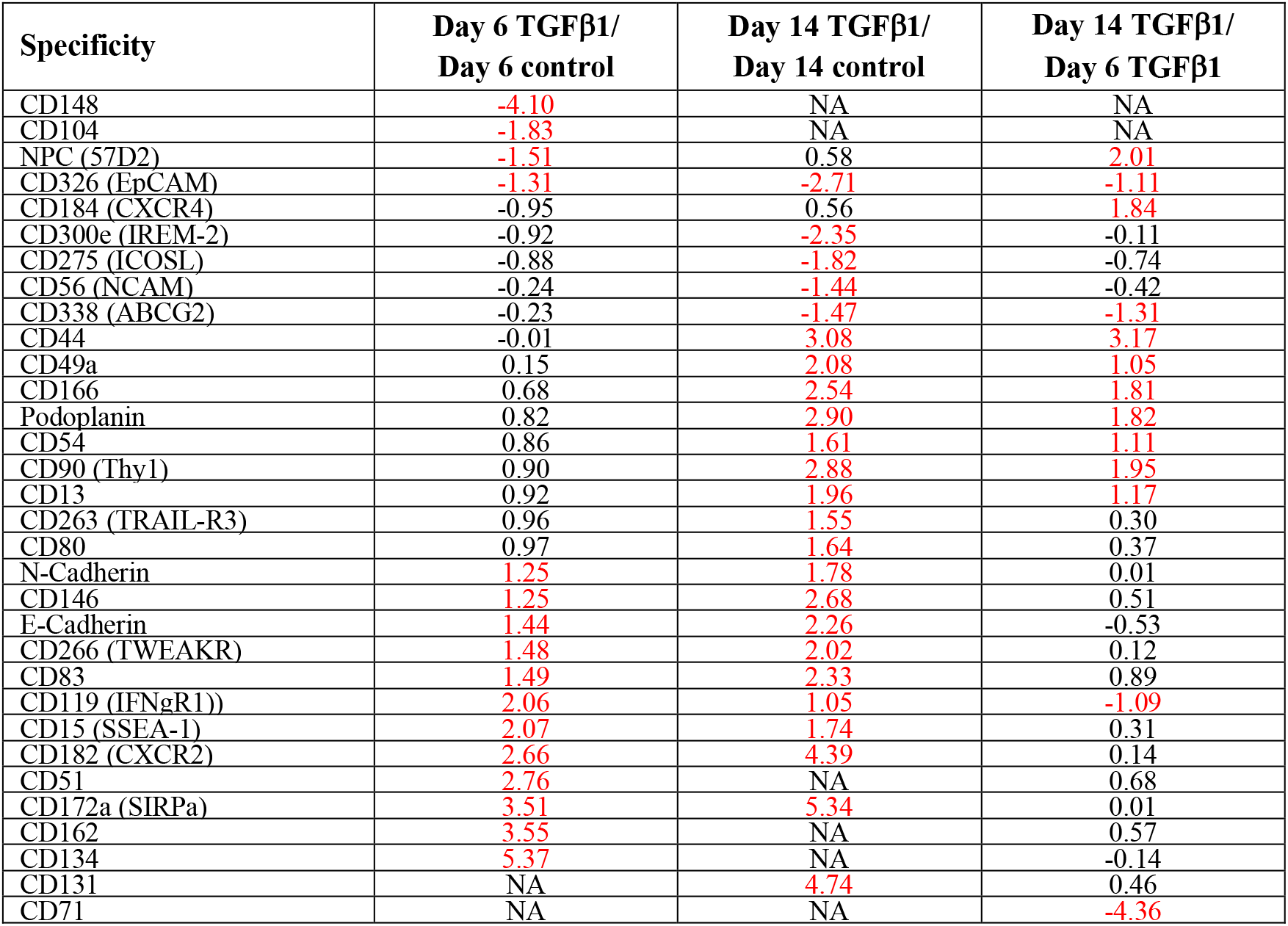
FACS screen results for HMLE cells showing log2 fold changes selected for at least two-fold differences (highlighted in red).

| Specificity | Day 6 TGFβ1/<br>Day 6 control | Day 14 TGFβ1/<br>Day 14 control | Day 14 TGFβ1/<br>Day 6 TGFβ1 |
| --- | --- | --- | --- |
| CD148 | -4.10 | NA | NA |
| CD104 | -1.83 | NA | NA |
| NPC (57D2) | -1.51 | 0.58 | 2.01 |
| CD326 (EpCAM) | -1.31 | -2.71 | -1.11 |
| CD184 (CXCR4) | -0.95 | 0.56 | 1.84 |
| CD300e (IREM-2) | -0.92 | -2.35 | -0.11 |
| CD275 (ICOSL) | -0.88 | -1.82 | -0.74 |
| CD56 (NCAM) | -0.24 | -1.44 | -0.42 |
| CD338 (ABCG2) | -0.23 | -1.47 | -1.31 |
| CD44 | -0.01 | 3.08 | 3.17 |
| CD49a | 0.15 | 2.08 | 1.05 |
| CD166 | 0.68 | 2.54 | 1.81 |
| Podoplanin | 0.82 | 2.90 | 1.82 |
| CD54 | 0.86 | 1.61 | 1.11 |
| CD90 (Thy1) | 0.90 | 2.88 | 1.95 |
| CD13 | 0.92 | 1.96 | 1.17 |
| CD263 (TRAIL-R3) | 0.96 | 1.55 | 0.30 |
| CD80 | 0.97 | 1.64 | 0.37 |
| N-Cadherin | 1.25 | 1.78 | 0.01 |
| CD146 | 1.25 | 2.68 | 0.51 |
| E-Cadherin | 1.44 | 2.26 | -0.53 |
| CD266 (TWEAKR) | 1.48 | 2.02 | 0.12 |
| CD83 | 1.49 | 2.33 | 0.89 |
| CD119 (IFNγR1) | 2.06 | 1.05 | -1.09 |
| CD15 (SSEA-1) | 2.07 | 1.74 | 0.31 |
| CD182 (CXCR2) | 2.66 | 4.39 | 0.14 |
| CD51 | 2.76 | NA | 0.68 |
| CD172a (SIRPa) | 3.51 | 5.34 | 0.01 |
| CD162 | 3.55 | NA | 0.57 |
| CD134 | 5.37 | NA | -0.14 |
| CD131 | NA | 4.74 | 0.46 |
| CD71 | NA | NA | -4.36 |

**Table 2:** FACS screen results for MCF10A cells showing log2 fold changes selected for at least two-fold differences (highlighted in red).

| Specificity | Day 2 TGFβ1/<br>Day 2 control | Day 4 TGFβ1/<br>Day 4 control | Day 8 TGFβ1/<br>Day 8 control |
| --- | --- | --- | --- |
| CD201 (EPCR) | -5.58 | NA | NA |
| CD148 | -3.46 | NA | 1.95 |
| CD165 | -2.23 | -2.51 | NA |
| E-Cadherin | -1.61 | -2.17 | -1.70 |
| MSC (W3D5) | -1.51 | -3.75 | -2.72 |
| Notch 1 | -1.38 | 0.97 | 0.53 |
| CD1a | -1.35 | 1.52 | -2.70 |
| CD9 | -1.28 | -2.22 | -1.61 |
| CD97 | -1.17 | -2.82 | -2.44 |
| CD111 | -0.99 | -2.00 | -2.65 |
| CD70 | -0.89 | -1.20 | -1.44 |
| CD298 | -0.87 | -1.92 | -1.89 |
| Notch 2 | -0.79 | -1.18 | -1.25 |
| CD55 | -0.77 | -1.14 | -1.30 |
| CD96 | -0.71 | 3.61 | 1.60 |
| CD325 | -0.62 | 1.57 | 1.44 |
| EGFR | -0.50 | -0.33 | -1.28 |
| CD56 (NCAM) | -0.45 | 0.27 | 2.45 |
| CD46 | -0.43 | -1.23 | -1.45 |
| TCR Vb8 | -0.36 | -1.48 | 1.85 |
| CD95 | -0.35 | -1.02 | -1.24 |
| CD11b (activated) | -0.34 | NA | 1.02 |
| CD338 (ABCG2) | -0.33 | 2.58 | 0.45 |
| MSC (W5C5) | -0.30 | -1.80 | -1.63 |
| Tim-4 | -0.25 | 1.54 | 0.46 |
| Siglec-10 | -0.22 | 1.05 | 0.50 |
| DR3 (TRAMP) | -0.22 | 0.47 | 1.20 |
| Siglec-9 | -0.20 | 1.03 | 0.46 |
| CD15 (SSEA-1) | -0.14 | 1.12 | 0.66 |
| Notch 3 | -0.12 | 2.35 | 0.78 |
| CD115 | -0.10 | 0.08 | 1.78 |
| b2-microglobulin | -0.09 | -0.57 | -1.11 |
| CD158a/h | -0.08 | -1.06 | 0.07 |
| CD255 (TWEAK) | -0.06 | 0.84 | 1.49 |
| CD156c (ADAM10) | 0.00 | -1.05 | -1.04 |
| CD47 | 0.02 | -0.45 | -1.05 |
| CD39 | 0.11 | -2.84 | 0.34 |
| CD49f | 0.13 | -0.83 | -1.07 |
| CD1d | 0.14 | 2.14 | 0.03 |
| Tim-1 | 0.15 | 1.14 | 0.52 |
| CD88 | 0.16 | 0.10 | -2.60 |
| CD215 (IL-15Ra) | 0.18 | 1.24 | 0.58 |
| HLA-E | 0.19 | 1.42 | 0.56 |
| CD86 | 0.20 | 3.31 | -0.16 |
| HLA-A2 | 0.24 | 1.38 | 0.08 |
| CD66a/c/e | 0.24 | -1.93 | -3.58 |
| CD24 | 0.26 | 0.02 | 1.11 |
| HER2 | 0.28 | 1.60 | NA |
| CD101 (BB27) | 0.31 | -1.00 | -0.63 |
| CD167a | 0.31 | 1.37 | 1.72 |
| IGF-1R | 0.34 | 0.65 | 1.11 |
| CD104 | 0.35 | -1.05 | -1.27 |
| CD89 | 0.35 | -2.28 | -3.49 |
| CD268 | 0.47 | 1.39 | -0.49 |
| Notch 4 | 0.47 | -0.93 | -1.97 |
| CD220 | 0.48 | 0.75 | -1.07 |
| CD252 (OX40L) | 0.48 | 1.08 | 1.37 |
| CD141 | 0.56 | -1.01 | 0.86 |
| CD318 (CDCP1) | 0.60 | -0.23 | -1.45 |
| CD63 | 0.64 | 1.07 | 1.23 |
| CD114 | 0.65 | 0.25 | 1.48 |
| CD83 | 0.76 | 1.54 | 1.46 |
| CD258 | 0.77 | 1.37 | -0.75 |
| CD105 | 0.87 | 1.14 | 1.77 |
| CD266 | 1.02 | 0.79 | 0.37 |
| CD80 | 1.04 | -0.18 | -0.54 |
| CD49a | 1.14 | 1.31 | 1.19 |
| TCR Vb23 | 1.15 | 1.55 | 0.62 |
| CD172a (SIRPa) | 1.22 | 0.96 | 0.57 |
| CD13 | 1.30 | 1.44 | 1.20 |
| CD116 | 1.36 | 1.36 | 0.43 |
| CD146 | 1.38 | 2.36 | 2.07 |
| CD5 | 1.49 | 0.00 | -0.02 |
| CD1b | 1.51 | -0.35 | 0.32 |
| CD138 | 1.76 | 0.26 | -0.55 |
| CD73 | 2.48 | 3.16 | 1.82 |
| CD131 | 2.54 | 0.19 | 0.73 |
| Podoplanin | 2.87 | 3.33 | 2.44 |
| CD51 | 2.95 | 2.96 | 2.25 |
| CD49d | 6.52 | 3.79 | 0.51 |
| CD273 | 7.98 | 2.69 | 4.51 |
| FcRL6 | NA | 0.04 | -1.16 |
| HLA-ABC | NA | NA | -2.26 |
| CD71 | NA | NA | -3.19 |

**Table 3:** FACS screen results for HTER cells showing log2 fold changes selected for at least two-fold differences (highlighted in red).

| Specificity | Day 8 4OHT/<br>Day 0 control | Day 12 4OHT/<br>Day 0 control | Day 12 4OHT/<br>Day 8 4OHT |
| --- | --- | --- | --- |
| CD20 | -4.17 | -0.57 | 3.60 |
| CD49d | -2.67 | -3.07 | -0.39 |
| CD300F | -2.36 | NA | NA |
| CD28 | -2.12 | -1.86 | 0.27 |
| CD201 (EPCR) | -1.99 | 0.44 | 2.43 |
| CD56 (NCAM) | -1.84 | -0.53 | 1.31 |
| CD70 | -1.43 | 0.32 | 1.75 |
| Notch 3 | -1.39 | -0.11 | 1.29 |
| CD24 | -1.33 | -1.23 | 0.10 |
| EpCAM | -1.32 | -3.72 | -2.40 |
| CD335 (NKP46) | -1.19 | -1.13 | 0.06 |
| CD1c | -0.93 | 0.39 | 1.32 |
| CD340 (HER2) | -0.88 | 0.70 | 1.59 |
| CD271 | -0.87 | 0.40 | 1.28 |
| CD85d (ILT4) | -0.86 | 0.65 | 1.51 |
| CD170 (Siglec-5) | -0.82 | -2.39 | -1.56 |
| CD71 | -0.80 | -1.40 | -0.60 |
| CD275 (ICOSL) | -0.79 | 1.00 | 1.78 |
| CD104 | -0.61 | 0.93 | 1.54 |
| CD109 | -0.57 | 0.86 | 1.43 |
| HLA-E | -0.40 | 1.09 | 1.48 |
| CD95 | -0.16 | 1.09 | 1.25 |
| CD221 (IGF-1R) | -0.06 | 1.10 | 1.16 |
| CD252 (OX40L) | 0.06 | 1.19 | 1.13 |
| CD119 (IFNgR1) | 0.07 | 1.09 | 1.02 |
| CD148 | 0.09 | 1.19 | 1.10 |
| CD33 | 0.20 | 1.47 | 1.27 |
| CD73 | 0.33 | 1.26 | 0.93 |
| MAIR-II | 0.35 | 1.17 | 0.82 |
| EGFR | 0.37 | 1.47 | 1.11 |
| CD83 | 0.43 | 1.53 | 1.10 |
| Tim-1 | 0.44 | 1.45 | 1.01 |
| CD79b | 0.46 | 1.09 | 0.63 |
| CD51 | 0.50 | 1.14 | 0.64 |
| HLA-A,B,C | 0.73 | 1.11 | 0.37 |
| CD44 | 0.86 | 1.17 | 0.30 |
| MSC (W5C5) | 0.90 | 2.75 | 1.86 |
| CD90 (Thy1) | 0.94 | 1.45 | 0.51 |
| CD200 (OX2) | 0.95 | 2.06 | 1.11 |
| CD255 (TWEAK) | 0.97 | 2.09 | 1.12 |
| CD93 | 1.11 | 1.83 | 0.72 |
| HLA-A2 | 1.16 | 1.16 | 0.00 |
| CD266 (TWEAK-R) | 1.31 | 0.07 | -1.24 |
| MSC (W3D5) | 1.56 | 3.75 | 2.19 |
| CD10 | 2.02 | 3.62 | 1.60 |
| CD38 | 5.61 | 4.69 | -0.92 |
| Notch 1 | NA | 2.44 | NA |
| CD290 | NA | NA | -2.71 |

**Table 4:**
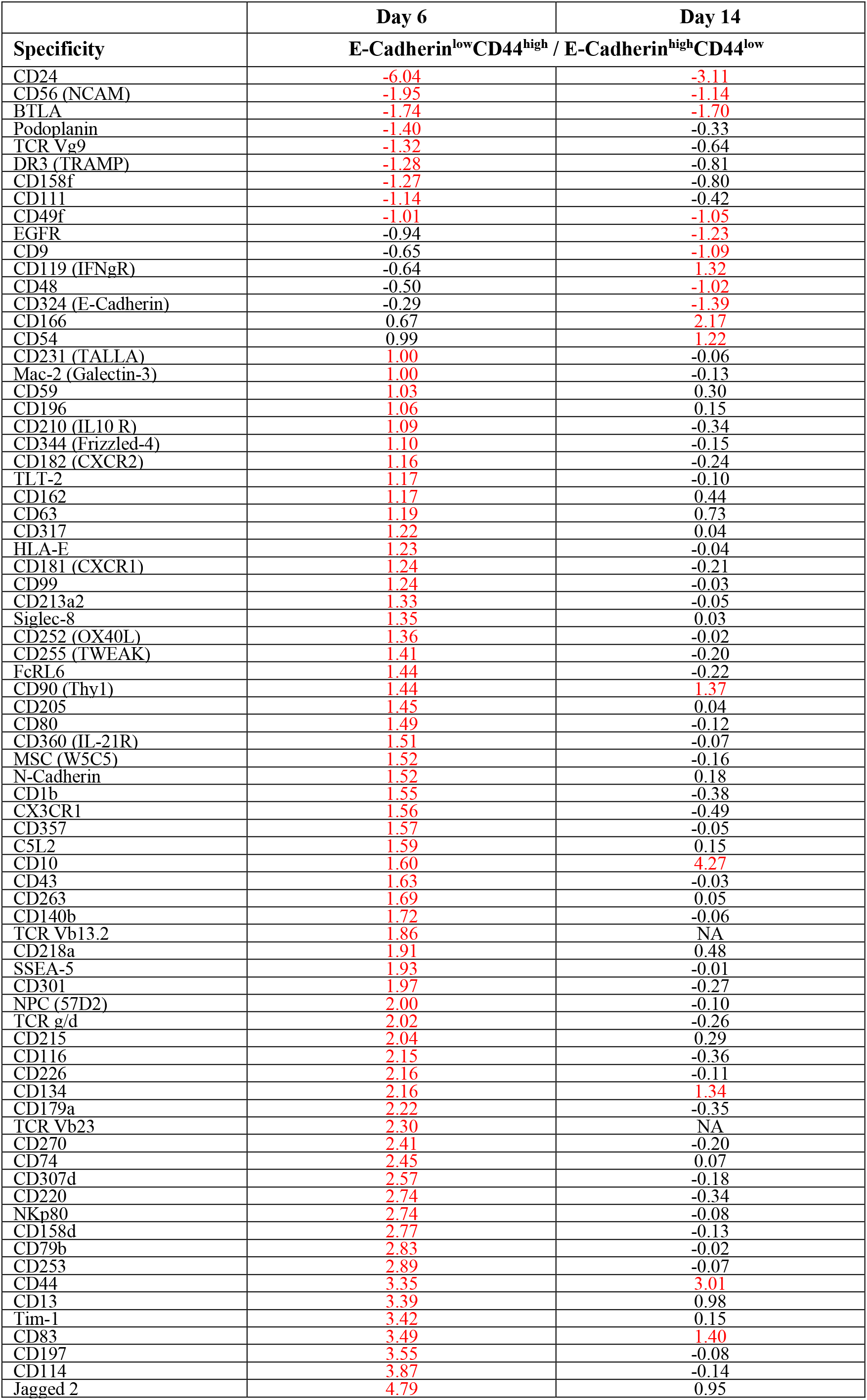
FACS screen results for TGFβ1-treated HMLE cells showing log2 fold changes selected for at least two-fold differences (highlighted in red).

**Table 5:** FACS screen results for TGFβ1-treated MCF10A cells showing log2 fold changes selected for at least two-fold differences (highlighted in red).

| Specificity | Day 2 | Day 4 | Day 8 |
| --- | --- | --- | --- |
|  | EpCAM<sup>low</sup> E-Cadherin<sup>low</sup> /<br>EpCAM<sup>high</sup> E-Cadherin<sup>high</sup> |  |  |
| EGFR | -2.30 | -1.70 | -2.49 |
| CD24 | -1.99 | -2.63 | -2.45 |
| CD56 (NCAM) | -1.89 | -1.93 | -1.31 |
| Notch 1 | -1.81 | -1.58 | -1.28 |
| CD9 | -1.63 | -2.44 | -2.75 |
| CD148 | -1.43 | -1.82 | -1.85 |
| DR3 (TRAMP) | -1.43 | -1.39 | -1.58 |
| Tim-1 | -1.30 | -2.03 | -2.69 |
| Pre-BCR | -1.25 | -1.87 | -2.42 |
| CD49f | -1.22 | -1.32 | -1.35 |
| CD140b | -1.17 | -1.58 | -3.00 |
| CD48 | -1.15 | -1.58 | -1.50 |
| CD131 | -1.14 | -1.27 | -1.66 |
| CD95 | -1.14 | -1.20 | -1.21 |
| Integrin a9b1 | -1.12 | -1.51 | -1.81 |
| TLT-2 | -1.09 | -1.65 | -1.75 |
| CD263 | -1.00 | -1.18 | -1.92 |
| CD82 | -1.00 | -1.07 | -1.29 |
| CD10 | 1.15 | 2.14 | 1.37 |

**Table 6:** FACS screen results for 4OHT-treated HTER cells showing log2 fold changes selected for at least two-fold differences (highlighted in red).

|  | Day 8 | Day 12 | Day 12 / Day 8 |
| --- | --- | --- | --- |
| Specificity | E-Cadherin<sup>low</sup> / E-Cadherin<sup>high</sup> |  |  |
| EpCAM | -4.19 | -5.93 | -6.55 |
| CD275 | -3.94 | -1.25 | 5.48 |
| CD220 | -3.18 | -2.10 | -2.15 |
| CD104 | -2.97 | -10.04 | -5.46 |
| CD271 | -2.58 | -2.93 | 1.03 |
| CD282 | -2.06 | NA | NA |
| CD33 | -1.76 | -1.18 | 1.88 |
| Notch 3 | -1.75 | 1.29 | 4.09 |
| CD24 | -1.57 | -2.29 | -1.27 |
| CD170 | -1.50 | -1.29 | -2.04 |
| CD15 | -1.37 | -1.23 | -0.10 |
| CD9 | -1.29 | -1.62 | 0.57 |
| CD201 | -1.28 | -0.49 | 3.31 |
| CD56 | -1.22 | -1.38 | 1.55 |
| CD324 | -1.20 | -1.42 | 0.44 |
| CD261 | -1.11 | -1.15 | 0.46 |
| CD338 | -1.10 | -1.76 | -1.57 |
| Tim-1 | -0.91 | -0.21 | 1.34 |
| CD11b | -0.88 | -1.04 | 0.17 |
| EGFR | -0.82 | -1.69 | 0.33 |
| CD141 | -0.80 | -0.86 | 0.55 |
| CD111 | -0.79 | -1.17 | 0.26 |
| CD340 | -0.74 | -0.13 | 2.02 |
| CD262 | -0.71 | -0.01 | 1.64 |
| CD52 | -0.69 | -2.86 | -1.56 |
| CD49f | -0.67 | -1.45 | -0.79 |
| MSC | -0.66 | -0.87 | 2.02 |
| CD221 | -0.64 | -0.62 | 1.14 |
| MSC | -0.62 | -0.89 | 1.43 |
| CD95 | -0.60 | -0.22 | 1.68 |
| CD34 | -0.58 | -0.37 | 1.16 |
| CD109 | -0.57 | -0.76 | 1.30 |
| CD318 | -0.57 | -1.23 | -1.03 |
| CD81 | -0.55 | 0.01 | 1.34 |
| BTLA | -0.54 | -1.07 | 0.40 |
| CD252 | -0.49 | -0.24 | 1.38 |
| CD119 | -0.48 | 0.94 | 1.84 |
| HLA-E | -0.48 | -0.39 | 1.59 |
| CD197 | -0.43 | -0.29 | 1.00 |
| CD266 | -0.41 | -0.62 | -1.49 |
| MAIR-II | -0.37 | -0.01 | 1.24 |
| CD94 | -0.33 | 0.07 | 1.40 |
| Mac-2 | -0.32 | -0.27 | 1.04 |
| CD277 | -0.14 | 0.16 | 1.36 |
| CD166 | -0.13 | 0.76 | 1.13 |
| CD200 | -0.02 | 0.20 | 1.13 |
| CD73 | 0.01 | 0.41 | 1.52 |
| CD70 | 0.07 | 1.33 | 2.84 |
| CD290 | 0.21 | NA | NA |
| CD83 | 0.24 | 1.08 | 1.68 |
| CD85d | 0.42 | -1.34 | 0.79 |
| CD304 | 0.51 | NA | -0.74 |
| CD10 | 0.93 | 2.30 | 2.45 |
| CD1c | 1.44 | -2.06 | -0.69 |
| CD44 | 2.04 | 3.41 | 0.55 |
| Siglec-9 | NA | 4.95 | 1.84 |
| CD314 | NA | 0.39 | 1.01 |

**Figure 3.**
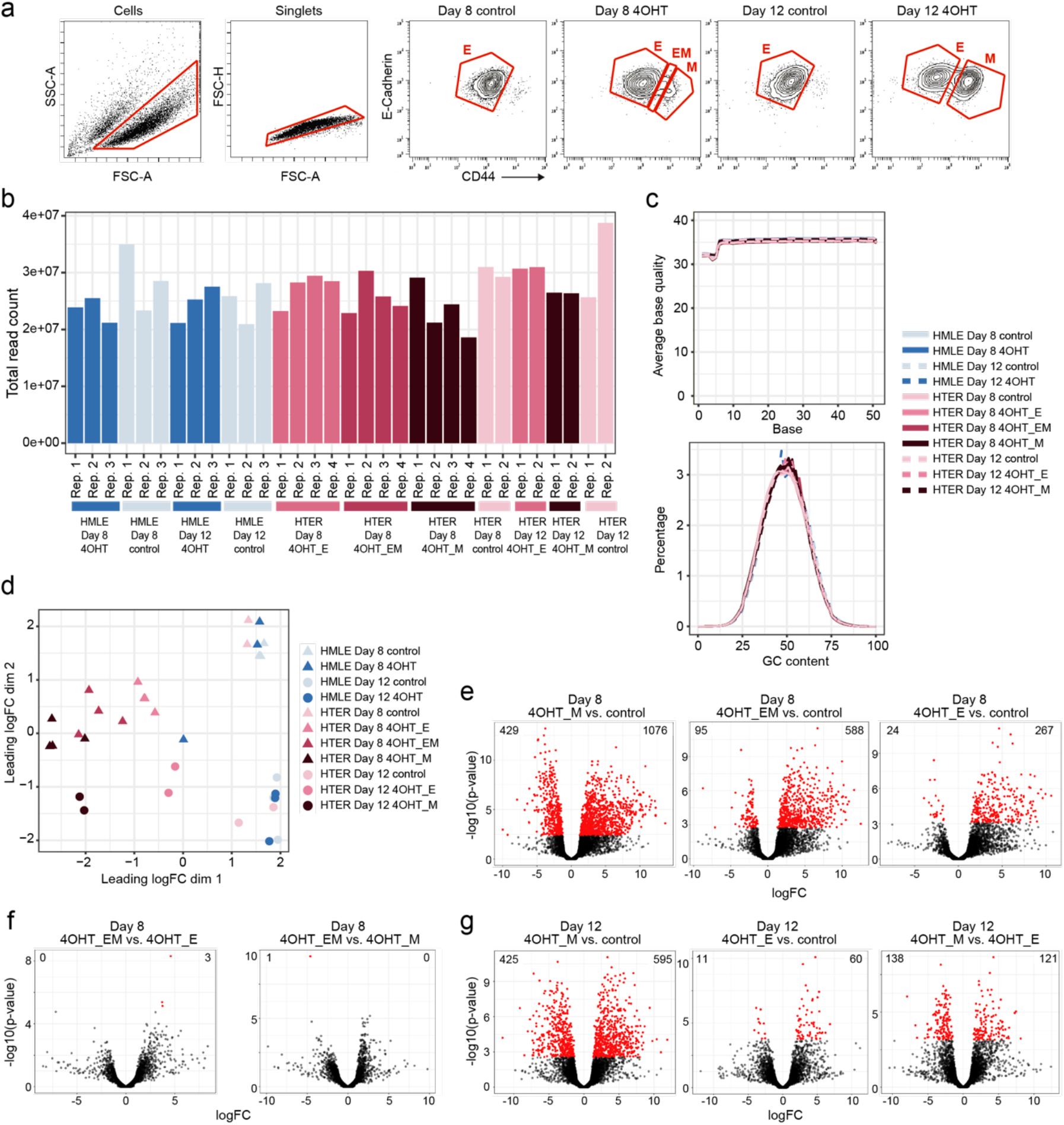
FACS surface protein profiling of EMT-undergoing mammary epithelial cells. (a) Gating to select populations with E-Cadherin^high^CD44^low^ or E-Cadherin^low^CD44^high^ phenotype. (b) Gating to select populations with E-Cadherin^high^EpCAM^high^ or E-Cadherin^low^EpCAM^low^ phenotype. (c) Gating to select populations with E-Cadherin^high^ or E-Cadherin^low^ phenotype. (d) Histogram overlays of HMLEs comparing CD90 levels in TGFβl-treated versus control (left panel) and CD44 levels in the E-Cadherin^high^CD44^low^ and E-Cadherin^low^CD44^high^ populations (right panel). (e) Proteins that were more than two-fold regulated between treated cells and control (upper panel) or between treated cells with E-Cadherin^high^ or E-Cadherin^low^ phenotype (lower panel). (f) Candidate antibody panel for mass cytometry analysis.

Based on these FACS screen results and the RNA sequencing analysis, we assembled a panel of candidate targets to assess phenotypic heterogeneity during EMT in more depth using a multiplex mass cytometry workflow (Figure 3f, Online-only Table 20).

### Mass cytometric profiling of EMT phenotypes

Mass cytometry is uniquely suited to assess phenotypic heterogeneity during EMT due to its ability to measure about 40 targets on the single-cell level^20,47^. To ensure high data quality, all antibodies against the candidate targets were titrated and validated using different cell lines and conditions (Figure 4a). We then selected EMT-undergoing and control samples at multiple time points for each of the HMLE, HTER, HSER, and MCF10A cell lines, totalling 92 samples (Figure 4b). The single-cell suspensions were fixed and mass-tag barcoded^24^ to allow the pooling and simultaneous antibody staining of the samples (Methods). We used an antibody against cleaved CASPASE-3 and cleaved poly(ADP-ribose)-polymerase 1 (PARP1) to exclude apoptotic cells, yielding more than 1 million live cells for downstream analysis (Figure 4c). Comparing three biological replicates of the MCF10A cell line using the dimensionality reduction algorithm Uniform Manifold Approximation and Projection (UMAP)^48^ showed a strong similarity of the triplicates and discrimination of treated and control samples, except for day 2 control cells (Figures 4d and 4e; Methods). Comparing the triplicates of the HMLE cell line using UMAP also confirmed a strong similarity, however, treated and control samples were less separable (Figures 4f and 4g). Applying the t-distributed stochastic neighbor embedding (t-SNE)^30^ dimensionality reduction algorithm to all samples visualized the phenotypic diversity of EMT-undergoing cells between the different cell lines and in comparison with the respective control (Figures 4h and 4i). In MCF10A cells, we observed a co-upregulation of CD44, Podoplanin, CD146, and CD51 upon EMT induction compared with control and concomitant downregulation of E-Cadherin and K5. In the HMLE, HTER, and HSER cell lines, Vimentin, CD44, CD90, CD51, and CD10 were co-upregulated in EMT-undergoing cells compared with control (Figures 4h and 4i). In conclusion, we assembled an antibody panel for multiplex mass cytometry characterization of EMT and discovered a vast phenotypic diversity of EMT states among four widely used human *in vitro* models of this process.

**Figure 4.**
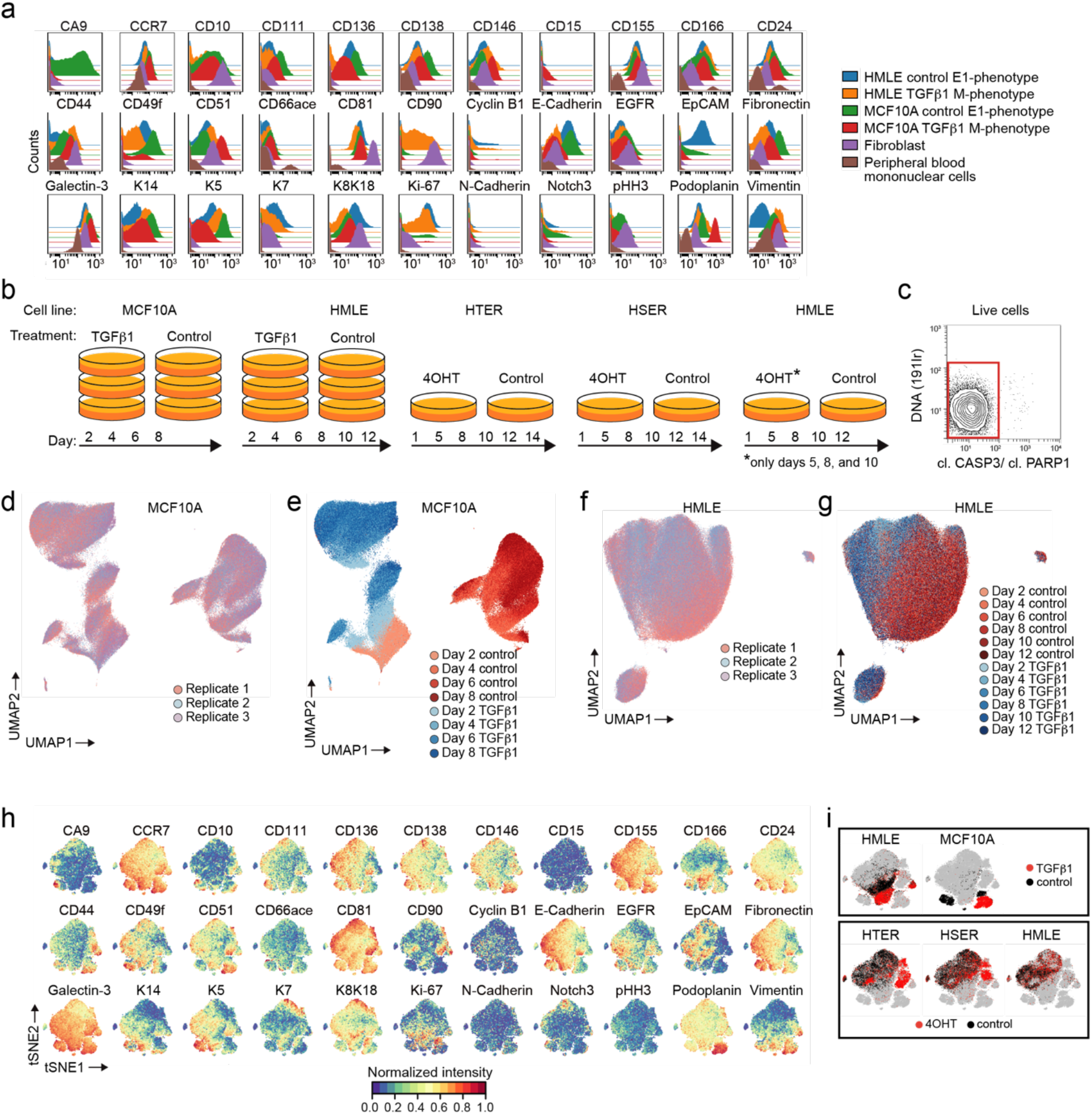
Multiplex mass cytometry profiling of EMT phenotypes. (a) Histogram overlays showing the antibody panel performance. (b) Types of samples collected for mass cytometry. (c) Gating to select live cells. (d-e) UMAPs showing TGFβ1-treated and control MCF10As colored by biological replicates (d) and by day and treatment (e). (f-g) UMAPs showing TGFβ1-treated and control HMLEs colored by biological replicates (f) and by day and treatment (g). (h) t-SNE maps showing the expression of markers on 52,000 cells after a 0 to 1 normalization. For each cell line, 1,000 representative cells were chosen from control and treated samples at all time points as indicated in (b). Only one replicate was used for MCF10As and HMLEs. (i) t-SNE maps as in (h), highlighting in black the cells from the indicated cell lines. For each line, both control and treated cells are shown.

## Code Availability

The code used for RNA sequencing data analysis can be found on GitHub^41^ and can be accessed without restrictions. Please refer to the Statistical Analysis section above for more details on software versions.

## Acknowledgements

We thank the Bodenmiller lab and the Robinson lab for fruitful discussions. We thank Stéphane Chevrier and the University of Zurich Cytometry Facility for advice and help regarding all FACS experiments. We thank Vito R. T. Zanotelli for advice regarding data visualizations. We thank the ETH Zurich Genomics Facility Basel for excellent RNA sequencing service. We thank the Weinberg lab for their gift of the HMLE, HMLE-Twist-ER, and HMLE-Snail-ER cell lines. B.B.’s research is funded by a SNSF R’Equip grant, a SNSF Assistant Professorship grant, the SystemsX Transfer Project “Friends and Foes”, the SystemsX MetastasiX and PhosphoNetX grants, and by the European Research Council (ERC) under the European Union’s Seventh Framework Program (FP/2007-2013)/ERC Grant Agreement n. 336921. M.D.R. acknowledges support from UZH’s URPP Evolution in Action.

## Author contributions

J.W. and B.B. conceived the study. J.W. performed the cell culture experiments together with M.M. and A.J.. J.W. and M.M. performed the FACS surface marker screens with help from A.J. and the mass cytometry stainings with the corresponding data processing and interpretation. J.W. performed the FACS sorting and RNA isolation experiments prior to RNA sequencing. C.S. and M.D.R. performed RNA sequencing data analysis. N.D. performed the UMAP data visualizations. J.W., N.d.S., and B.B. wrote the manuscript with input from all authors.

